# Multi-Atlas Image Soft Segmentation via Computation of the Expected Label Value

**DOI:** 10.1101/2020.10.08.331553

**Authors:** Iman Aganj, Bruce Fischl

## Abstract

The use of multiple atlases is common in medical image segmentation. This typically requires deformable registration of the atlases (or the average atlas) to the new image, which is computationally expensive and susceptible to entrapment in local optima. We propose to instead consider the probability of all possible atlas-to-image transformations and compute the *expected label value (ELV)*, thereby not relying merely on the transformation deemed “optimal” by the registration method. Moreover, we do so without actually performing deformable registration, thus avoiding the associated computational costs. We evaluate our ELV computation approach by applying it to brain, liver, and pancreas segmentation on datasets of magnetic resonance and computed tomography images.

## I. Introduction

AUTOMATIC image segmentation is often a central step in medical imaging studies, enabling the analysis of specific regions of interest (ROIs). In supervised segmentation, an algorithm segments a new image using the information derived from a training dataset of images that are accompanied with gold-standard (e.g. manually delineated) ROI labels. Two popular approaches to supervised image segmentation use multiple atlases [1-4] and deep neural networks [5, 6]. In multiatlas-based segmentation of a new image, atlas images are (or a mean template image is) deformably registered to the new (to-be-segmented) image. The manual labels are then propagated into the new image space using the computed transformations, and fused to form the new labels.

Deformable registration of the atlas images to the new image is computationally very demanding (except for recent deeplearning based approaches [7-10]) and is the bottleneck of atlas-based segmentation. To improve computational efficiency, it has been proposed to use only a subset of atlases [11], albeit at the price of discarding a portion of the available training data.

The transformation resulting from registration guides label propagation from the atlas to the new image. Being an iterative non-convex optimization, image registration is prone to becoming trapped in local optima, potentially leading to inaccurate propagation of the labels. Moreover, different but equally reasonable transformations may produce similar values for the registration objective function (within its margin of error). Thus, even if the global optimum is found, choosing it as the single correct transformation would disregard valuable information provided by other potentially valid transformations. Such a globally optimal solution is also rarely robust, as it is sensitive to disturbances of or changes to input images, or variations in acquisition parameters. To alleviate this issue, uncertainty in registration has been incorporated into Bayesian segmentation by approximating the marginalization over registration parameters via Markov Chain Monte Carlo techniques [12], which, even though efficiently implemented, would further increase the computational costs. Local measures of uncertainty in deformable registration have also been used to improve the sensitivity of the label propagation in atlas-based segmentation [13, 14].

In this work, we present a new atlas-based method for soft (i.e., fuzzy or probabilistic) segmentation of images, which – instead of attempting to determine a single correct label – produces the expected value of the label at each voxel of the new image, while considering the probability of possible atlas-to-image transformations. This is accomplished without either explicitly sampling from the transformation distribution (which would be intractable) or running the costly deformable registration in training or testing stages. We create a single image from the training data, which we call the *key*. Then, for a new image (after affine alignment, if necessary), we compute the *expected label value (ELV)* map simply via a convolution with the key, which is efficiently performed using the fast Fourier transform (FFT). Our fuzzy ELV map is therefore a robust combination of labels suggested by atlas-to-image transformations, weighted by a measure of the transformation validity. This soft segmentation can be further used to initiate a subsequent hard-segmentation procedure. We validate our approach through brain, liver, and pancreas segmentation experiments on magnetic resonance (MR) and computed tomography (CT) images.

This article extends our preliminary conference version [15]. In particular, we have improved the method as well as expanded our empirical evaluation by including several new datasets. Moreover, our Matlab toolbox is now publicly available (https://www.nitrc.org/projects/elv). In the following, we describe the proposed method in detail (Section II and the appendices) and present experimental results (Section III) along with some concluding remarks (Section IV).

## II. Methods

### A. Segmentation from a Single Atlas

Let: ℝ^*d*^ → ℝ be the *d*-dimensional image to be segmented, and *J*: ℝ^*d*^ → ℝ an atlas image with the same contrast as, for which the manual label of a specific ROI has been provided as ^ℝ^ *L*: ℝ^*d*^ → 0,1 .^1^ For the new image, we wish to compute the expected value of the ROI label, *E*: ℝ^*d*^ → [0,1], which is a measure of likelihood of each voxel belonging to the ROI.

In traditional atlas-based image segmentation, the label *L* is propagated into the space of as *L* ° ***T***^*∗*^ (*I,J*),, where the transformation ***T***^*∗*^, is computed via registration as ***T***: ℝ^*d*^ → ℝ^*d*^ that maximizes the similarity between *I* and *J* ° ***T***.^2^ Here, instead, we propose to compute the *expected value* of the propagated *L*, while considering a probability for each possible transformation in 𝕋 ≔ ***T***: ℝ^*d*^ → ℝ^*d*^, as follows:

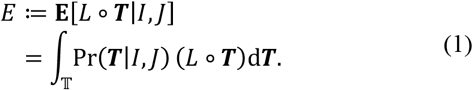

Equation (1) computes the ELV as an integral over the space of all transformations, which could be regarded as multiple (theoretically an infinite number of nested) integrals over the space of parameters representing ***T***. For free-form deformation, as considered here, Eq. (1) in fact includes a *d*-dimensional integral – with respect to the value of ***T*** – for each ∈ ℝ^*d*^. In standard atlas-based segmentation, Pr ***T***|*I,J*), is considered a Dirac delta, ∂ (***T*** − ***T***^*∗*^ (*I,J*),), whereas here we will consider a full probability distribution for it.

Using Bayes’ theorem, we can write the probability of the transformation given both the new and atlas images as:

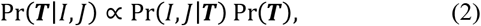

where the two right-hand-side factors correspond to the image similarity and the transformation regularity, respectively. For the former, we opt to use the inner product of the image and the transformed atlas, since it is expected to be higher when the two images are well aligned:

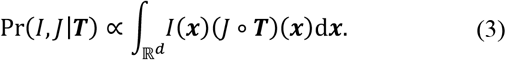

It is, however, well established that the inner product reflects the degree of alignment more effectively when only the *phase* information of the image is included [16, 17], which is how in practice we will proceed, as described in Section II.C. A discussion on our choice of the inner product of phase images as image similarity is provided in Appendix A. In the following, we first consider the case where ***T*** is only a translation.

#### 1) Translation

For a translation, ***T***(***x***)= (***x***)−**Δ**, the inner product in Eq. (3) becomes the cross-correlation of the image and the atlas, which is commonly used for image alignment [16, 17]:

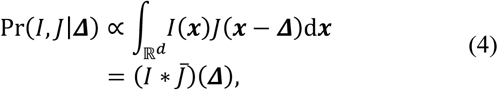

where *∗* denotes the convolution operator,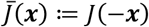. By assuming a flat prior for the shift (i.e., a constant Pr(**Δ**) and combining Eqs. (1), (2), and (4), the ELV at voxel y will be:

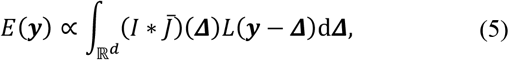

or,

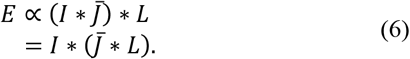

In the second line, we exploited the associativity property of convolution, which leads to the following expression for the ELV:

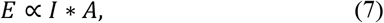

where we define and pre-compute the *key, A*, from the atlas, as:

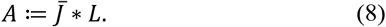

This convolution operation blurs the flipped atlas image and shifts it in the opposite direction of the ROI, thereby resulting in a key image, *A*, with the ROI roughly at its center.

Next, we will incorporate deformations in our transformation model.

#### 2) Deformation

To generalize the transformation ***T*** to include deformations, we will use the common Tikhonov prior on the regularity of the deformation field as the probability of the transformation:

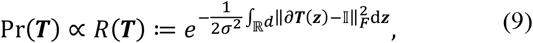

where ∂***T*** is the Jacobian matrix of ***T***, 𝕀 is the *d* × *d* identity matrix, and the constant parameter *A* represents a prior on the magnitude of the deformations. In Appendix B, we show that the ELV is still computed following Eq. (7), where the key, *A*, is initially computed as in Eq. (8), but then updated to incorporate the deformation. We show that we can approximate this update by an inhomogeneous blurring of the key, as:

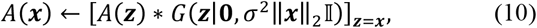

where *G*(· |***μ***, *Σ*) represents the Gaussian function with the mean ***μ*** and the co-variance matrix *Σ*. One can see that the size of the blurring kernel increases with the square root of the Euclidean distance from the center of *A* – i.e., the region corresponding to the label ROI (see Section II.A.1). Blurring a region in *A* decreases its contribution to soft segmentation by removing its high-frequency components prior to the convolution in Eq. (7). This means that Eq. (10) takes local deformations into account by giving a smaller weighting to regions in the atlas image that are farther from the ROI, making the information in such far areas less important.

Note that, even without the inhomogeneous blurring (i.e., *σ* = 0), the proposed method still encompasses – not just a single optimal, but – all possible translations, each partially contributing to the ELV with a weighting proportional to its degree of aligning. Given that a deformation field itself can be construed as many local translations, each of which would partially contribute to the ELV, one can see that, in some way, deformations are inherently accounted for.

The proposed model accounts for large translations, as well as local deformations, even though we do not run any deformable registration. As for rotation and global scaling, accounting for local deformations covers a small amount of them, and to allow for large amounts, we can initially affinely align the image and the atlas.

### B. Multiple Atlases

In case *N* atlases (affinely aligned in the same space) with manual labels are available, we will write Eq. (1) in the same fashion, as:

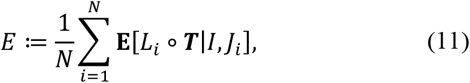

where *j*_*i*_ and *L*_*i*_ are the *i*^*th*^ pair of atlas and manual-label images, respectively. This will yield similar results as in Eqs. (7) and (10), with the only difference being Eq. (8), now generalized as:

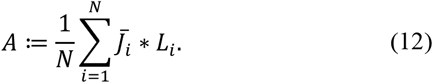

Note that even in the case of multiple atlases, the key, *A*, is a *single* image that is pre-computed from the training data.

Given that the to-be-segmented ROI is roughly at the center of the key, the regions whose relative positions with respect to the ROI are consistent across atlases are amplified in the key through constructive addition in Eq. (12), whereas those with relative positions varying from atlas to atlas due to inter-atlas deformations are washed out. Therefore, the relative weighting of the regions that are most informative for segmenting the desired ROI is elevated in the key thanks to aggregation of information from multiple atlases.

### C. Implementation

#### 1) Computation in the Fourier Domain

To create the key, *A*, we first ensure that the *N* training images are represented roughly in the same space; and if not, we affinely align them. By applying the convolution theorem to Eq. (12), we will then use FFT to initialize *A*:

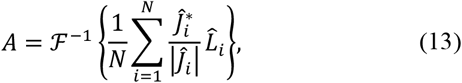

where the *hat* (^) sign and ℱ^-1^ represent the Fourier and inverse Fourier transforms, respectively, and *ĵ*^∗^ is the complex conjugate of *ĵ*. By only keeping the phase information of the image (i.e., normalizing *ĵ*_*i*_ by its magnitude), we create a sharper probability distribution for the aligning transformation in Eq. (3) [16, 17] (see Appendix A). In addition, this has an intensity normalization effect, preventing *A* from giving a different weighting to an atlas image due to its global intensity scaling. Next, for inhomogeneous blurring of the key (i.e., if *σ* > 0), we update *A* voxel-wise following Eq. (10) by multiplying and summing it with a varying discretized Gaussian kernel.

To segment a new image, *I*, we first make sure that it is correctly represented in the atlas space (otherwise, affinely align it to the mean atlas image using any existing affine registration tool), and then compute the ELV map from Eq. (7) as follows:

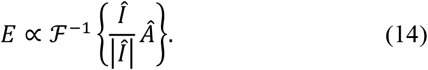

Note that *Â* is pre-computed from the atlases and kept offline. Different keys can be independently created for different organs for the purpose of multi-organ segmentation, in which case label overlap can be avoided using the soft-max operator [6] on the normalized probability maps of different organs, and via other label-fusion strategies [3, 4] such as STAPLE [18].

For hard segmentation of the organ (or structure) from the map, we threshold the map to keep a voxel subset with the volume 14% larger than that of an average organ (estimated from the atlases); see Appendix C for the rationale behind this choice.^3^ We then refine the mask by keeping the largest connected component (CC), as well as the CCs with at least half the volume of the largest CC, and then filling the holes.^4^

### 2) Second Pass

Once the initial ELV map is obtained, it can be refined by recalculating Eq. (14) while this time prioritizing the initial soft-segmented area. In our experiments, for instance, we used weighted versions of *A*(***x***) and *I*(x), as *A*(***x***) *G* (***x***|**0**, *s*^2^ 𝕀) and *I*(x) [*E* (***y***)*∗ G*(***y*** |**0**, *s*^2^𝕀)]_***y***= ***x***_, respectively, where the size of the Gaussian window (2*s*) was chosen to be roughly that of an average organ.

### 3) Intensity Prior

Given that using the phase image discards some image intensity information, one can further augment the computed ELV volume with image intensities. At a given voxel, the Bayes formula implies:

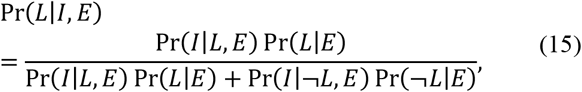

where *L* indicates that the given voxel belongs to the label, with ¬ the negation operator, and and *E* are the values of the image intensity and the computed ELV at the voxel, respectively. Were it known whether the voxel is included in the label or not, the image intensity would be conditionally independent of the ELV; i.e. Pr(*I* |*L, E*) = Pr (*I* |*L*)and Pr (*I* |¬*L, E*) = Pr(*I* |¬*L*). Using the ELV for Pr(*L*|*E*) then leads to:

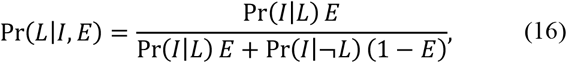

where Pr(*I* |*L*)and Pr(*I* |¬*L*)can be approximated by Gaussian functions of the intensity values of, with their parameters estimated from the atlases (or the image itself using the initial ELV map). We use the acronym ELV+IP to refer to this ELV map that has been modulated by the intensity prior (IP). For *E* to exhibit the properties of a probability, the ELV map needs to be normalized by its maximum, with any negative values projected to zero. Note that *E* was created using the *phase* images that had lost some image intensity information, meaning that the full image (*I*) in Pr (*L*| *I, E*) brings about new information not already included in *E*.

Pr(*L*| *I, E*)can even substitute for itself in the computation of the ELV, as – depending on the image contrast – it may better highlight the organ of interest, which is the most informative part of the image for segmentation. In that case, since the ELV map has not yet been computed, we use a constant *E* in Eq. (16) equal to the label-to-image volume ratio estimated from the atlases.

Several other post-processing steps are possible after this soft segmentation [1]. If binary segmentation is desired, the ELV map can then be thresholded (see Appendix C and Section II.C.1) or used as a seed region to subsequently initialize an unsupervised hard segmentation algorithm [15, 20].

## III. RESULTSand Discussion

We evaluated our ELV computation method on several medical image databases via leave-one-out cross validation. We inspected the images in each database to make sure that they were represented with the correct orientation in the same space, i.e. they were not misaligned due to dimension permutation or mirroring. Since we did not subsequently see any remaining global scaling and rotation that exceeded the inherent inter-subject variability, we did not affinely register the images.^5^ For each test image in a database, we created the key from the remainder of the images (i.e., labeled atlases) in the database following Eq. (12), computed the label for the test image, and report the Dice overlap coefficient between the computed label and the known label. Since optimizing for σ in Eq. (10) improved the Dice scores only negligibly (< 1%) in our initial benchmarking, we report our results in this section for the simple case with *σ* = 0.

As described in Section II.C, we computed the ELV map in two passes, modulated the ELV with the intensity prior in Eq. (15), where we used the atlases to estimate the means (except for the liver; see Section III.B) and standard deviations, and then hard-thresholded the probability maps to create masks. The algorithm chose a single CC from the mask in all experiments, except for 3% and 0.4% of the pancreas and hippocampus segmentation cases, respectively, where two CCs were chosen.

For data homogeneity in the abdominal segmentation experiments, we devised some inclusion criteria for each dataset, which were roughly chosen to minimize the number of excluded subjects (see below). Additional steps to preprocess the abdominal CT images included: smoothing the borders of each image, automatically removing the patient table (via thresholding the image and removing the lower-most one or two CCs), and using the intensity-prior image (Section II.C.3) instead of the abdominal image itself for ELV computation (thereby highlighting the organ amongst all other parts of the image).

### A. Brain

We first assessed the ability of our method to imitate FreeSurfer [21] in segmenting brain subcortical structures. We used T_1_-weighted MR images of 1224 subjects from the third release in the *Open Access Series of Imaging Studies (OASIS-3)* [22], normalized to the size 256×256×256 with 1 mm^3^ isotropic resolution. We considered the FreeSurfer-generated labels for 12 subcortical structures (left and right thalamus, caudate, putamen, pallidum, hippocampus, and amygdala) as “silver” standard and tried to reproduce the segmentation for each image via the proposed ELV approach. The median, mean, and standard error of the mean (SEM) of the resulting cross-validation Dice scores between the labels generated by ELV+IP and FreeSurfer are shown in Table I. Overall, the Dice score had a median of 0.782 and a mean of 0.766 ± (SEM) 0.001 across subjects and structures. (For the raw ELV, the median was 0.740 and the mean was 0.721 ± 0.001.) Since no manually delineated labels were used as the gold standard in this experiment, the results merely reveal how faithful the proposed approach is in reproducing FreeSurfer labels, rather than comparing the two methods.

**Table I.** Dice coefficients between ELV+IP and FreeSurfer labels in brain

| Structure | Hemisphere | Median | Mean | SEM |
| --- | --- | --- | --- | --- |
| Thalamus | Left | 0.751 | 0.744 | 0.001 |
|  | Right | 0.771 | 0.764 | 0.001 |
| Caudate | Left | 0.804 | 0.793 | 0.002 |
|  | Right | 0.807 | 0.796 | 0.002 |
| Putamen | Left | 0.808 | 0.797 | 0.002 |
|  | Right | 0.816 | 0.802 | 0.002 |
| Pallidum | Left | 0.758 | 0.685 | 0.005 |
|  | Right | 0.769 | 0.743 | 0.003 |
| Hippocampus | Left | 0.793 | 0.784 | 0.001 |
|  | Right | 0.806 | 0.799 | 0.001 |
| Amygdala | Left | 0.754 | 0.742 | 0.002 |
|  | Right | 0.758 | 0.748 | 0.002 |
| **All** | **Both** | **0.782** | **0.766** | **0.001** |

We then externally evaluated our approach by applying the key generated from the OASIS-3 images and FreeSurfer labels to segment two previously-unseen datasets of T_1_-weighted brain MR images with manual labels. The first dataset, the Internet Brain Segmentation Repository (IBSR) [23], included images from 18 subjects with the slice thickness of 1.5 mm and in-plane pixel sizes ranging from 0.84 mm to 1 mm, which we resampled to 1 mm^3^ isotropic voxel size. The second dataset [24] included 1 mm^3^ isotropic-voxel images from 39 subjects (28 healthy and 11 with dementia). We ran our ELV segmentation pipeline on each dataset once (without any further finetuning), resulting in median Dice scores of 0.69 (mean: 0.68 ± 0.01) and 0.677 (mean: 0.668 ± 0.004) for the two datasets, respectively. Note that the segmentation error in these results is the aggregate error from both the silver-standard labels used in training and the subsequent ELV segmentation. Further modulation with the intensity prior here deteriorated the results (median ELV+IP Dice: 0.56 and 0.624, respectively); this was not surprising, as the parameters of the image intensity prior had been estimated from another dataset (OASIS-3) with different image acquisition protocols, and the new images were not intensity-harmonized. The ELV itself, however, uses the phase images and is thus not so sensitive to inter-dataset variation in image intensity.

For comparison, in a similar experiment [25], a U-Net type convolutional neural network (CNN) was trained on 581 FreeSurfer-segmented T_1_-weighted brain images. The authors’ trained model produced mean Dice scores of 0.74 and 0.71 on two manually labeled test datasets. (The authors, however, did not compare the labels that they computed with FreeSurfer-generated labels.)

### B. Liver

Next, we used the training dataset of the public *Liver Tumor Segmentation (LiTS) Challenge* [26], which includes abdominal CT images with manually delineated labels for the normal tissue and lesions in the liver, provided by various clinical sites.

Although the subjects were scanned with contrast enhancement for liver lesions, we considered the entire (healthy and lesion) organ label in our experiments. 85 subjects passed our inclusion criteria, mainly the slice thickness being included in the header and no larger than 2 mm. The images were resampled in the space of the first image (where the Dice scores are reported) to (1.6mm)^3^ isotropic resolution, so they were all of the size 248×248×323.^6^

We computed the ELV map for each subject, an example of which is illustrated in Fig. 1 (left) for the representative subject (corresponding to the median final Dice score; see below). To create the ELV+IP map, we estimated the standard deviation of the intensities of the liver and the background from the 84 atlas subjects, using the manual labels and their dilated versions (by a sphere of radius 50), respectively. For stability, we estimated the *mean* intensity using the initial ELV mask of the test subject, given that lesion size and intensity varied from subject to subject. Next, we modulated the ELV with the intensity prior (Fig. 1, middle), created a new mask, and further refined it with an updated intensity mean estimated from this mask. For mask preparation, we also performed morphological *opening* with a spherical structuring element with the radius of 2 voxels while keeping the largest CC (i.e., eroding + keeping + dilating), which removed unwanted smaller structures attached to this relatively large ROI.

**Fig 1.**
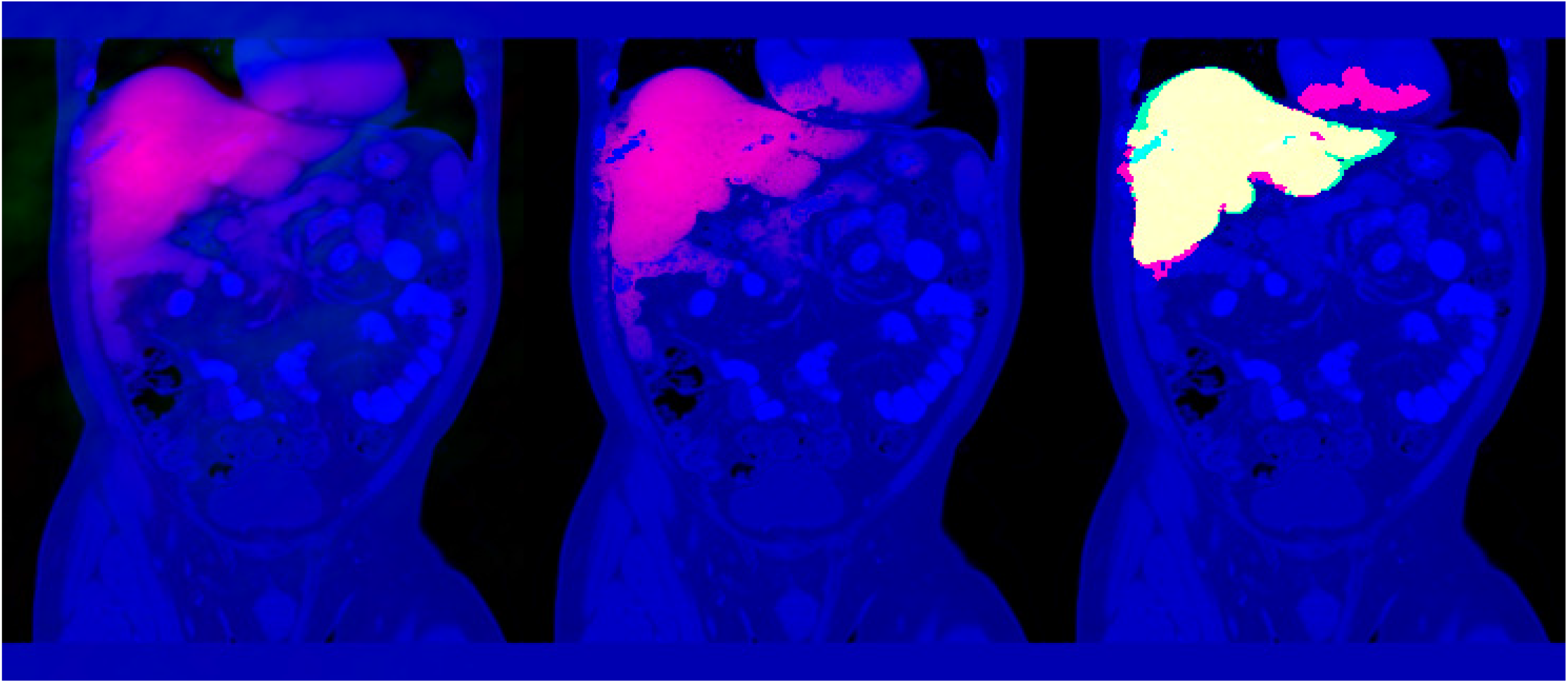
CT image (blue) of the representative subject (i.e., with median segmentation Dice score) in the LiTS dataset. The slice with the largest cross section with the manual label is shown. *Left:* The ELV map of the liver (red; occasional negative values in green). *Middle:* The ELV+IP map (red). *Right:* The resulting binary segmentation (red), the manual label (green), and their overlap (yellow). Intensities have been scaled for better visualization. A 3D video of this figure is available in the supplementary materials.

The cross-validation Dice coefficients between the masks computed from ELV+IP and manual labels (Fig. 1, right) had a median of 0.92 across subjects (mean: 0.91 ± 0.01). Dice scores, false positive rates (FPR), and false negative rates (FNR) are shown in Table II for the ELV and ELV+IP approaches, and a volumetric video of Fig. 1, as well as a video of ELV slices for all subjects are available in the supplementary materials. Among those results with lower (entire-organ) Dice scores, lesion regions were frequently the culprit, as the intensity prior, although generally improving the segmentation, partially excluded some of those regions. The relative portion of the lesion in the liver was significantly negatively correlated with the Dice score generated by the ELV+IP (r = -0.48, p < 10^−5^), but not the ELV (r = -0.01, p = 0.93). Repeating this experiment with *σ* = 0.1 resulted in a negligible change (< 0.0001) in the mean Dice score, which was not statistically significant (paired *t*-test p = 0.16).

**Table II.** Overlap of the automatic and manual liver labels

| Method | Dice Score | | FPR | FNR | Runtime (min): Mean ± SEM | |
| --- | --- | --- | --- | --- | --- | --- |
| | Median | Mean ± SEM | Mean ± SEM | | Training | Testing |
| ELV | 0.87 | $0.85 \pm 0.01$ | $0.46\% \pm 0.03\%$ | $8\% \pm 1\%$ | $31.7 \pm 0.2$ | $0.90 \pm 0.01$ |
| ELV+IP | 0.92 | $0.91 \pm 0.01$ | $0.19\% \pm 0.02\%$ | $9\% \pm 1\%$ | | $1.08 \pm 0.01$ |
| SyN deformable: Majority voting | 0.80 | $0.72 \pm 0.02$ | $0.13\% \pm 0.01\%$ | $37\% \pm 2\%$ | $300 \pm 4$ | |
| SyN deformable: Joint label fusion | 0.88 | $0.82 \pm 0.02$ | $0.07\% \pm 0.01\%$ | $26\% \pm 2\%$ | $375 \pm 4$ | |
| U-Net | 0.90 | $0.90 \pm 0.01$ | $0.26\% \pm 0.02\%$ | $8\% \pm 1\%$ | (GPU)<br>$131.0 \pm 0.2$ | (GPU)<br>$12.05 \pm 0.01$ |
| LiTS leaderboard | - | $0.84 \sim 0.97$ | | | - | |

To compare our method with state-of-the-art multi-atlas based image segmentation, we then used the `antsJointLabelFusion` function of the Advanced Normalization Tools (ANTs) [27] to segment the liver using both majority voting and joint label fusion [4]. For each test subject, the remaining 84 atlas images were registered to the test image, into whose space labels were then propagated and fused. We chose the *rigid + affine + deformable SyN* as the transformation type. As shown in Table II, joint label fusion (the best of the two) achieved a median Dice score of 0.88 (mean: 0.82 ± 0.02).

Next, we trained a 3D CNN with the U-Net architecture [6] for liver segmentation, which had 2 downsampling layers and 16 initial filters (at the first convolutional layer), and used the Dice coefficient as the objective function. We trained the network for 5 epochs using 3D sample patches of size 132×132×132 with a mini-batch size of 8. Through 10-fold cross-validation, each subject’s label was predicted by averaging the label scores of overlapping patches (10 voxels apart), binarizing the label, and keeping the largest CCs as we did for the ELV (which improved the results). As shown in Table II, the median Dice coefficient was 0.90 (mean: 0.90 ± 0.01). Increasing the number of initial filters to 32 at the price of a smaller patch size (100×100×100) lowered the Dice values.

For comparison, at the time of the submission of this article, the LiTS challenge website [26] reported mean Dice values for the liver on their test data ranging from 0.84 to 0.97 (disregarding the outlier results with mean Dice ≤ 0.35), with many of the methods applying deep learning.

To evaluate the runtimes, we ran the atlas-based methods on a compute cluster, allocating 4 CPU cores, 48GB of memory, and no graphics processing unit (GPU) to each computational job. The ELV method exploited the available 4 cores through Matlab’s internal parallelization of FFT, and ANTs did so via parallelization of atlas-to-image registrations (due to our use of the `-c 2 -j 4` options). Each U-Net experiment was run on an Nvidia RTX8000 GPU with 48GB of memory. The runtimes of all methods are shown in Table II. For the ELV+IP, the mean entire-experiment runtime for a test subject was 32.8 ± 0.2 min, of which 31.7 ± 0.2 min were spent in the training step (key creation). Soft segmentation from the key and subsequent hard segmentation, on the other hand, took only 0.86 ± 0.01 min and 0.229 ± 0.001 min, respectively. In contrast, the majority voting and joint label fusion methods of ANTs took 300 ± 4 min and 375 ± 4 min per test subject, respectively. The U-Net took 131.0 ± 0.2 min for training and 12.05 ± 0.01 min to segment a test subject. Given that the U-Net experiment was accelerated via a GPU, its runtime is not meant to be directly compared to those of the atlas-based methods that did not use GPUs.

### C. Pancreas

Lastly, we took a similar approach as in the previous subsection to segment the pancreas in two experiments, using two CT databases from *The Cancer Imaging Archive (TCIA)* acquired at the National Institutes of Health (NIH) Clinical Center [28, 29] (82 subjects) and from the *Memorial Sloan Kettering Cancer Center* [30] (225 subjects; those with slice thickness of 2∼3 mm). The labels created from the ELV+IP map in cross-validation had a median Dice score of 0.59 (mean: 0.56 ± 0.02) for the former database and a median Dice score of 0.50 (mean: 0.48 ± 0.01) for the latter database. (For the raw ELV, the median was 0.57 and 0.49, respectively.) Note that the pancreas in the second dataset included lesions (although no significant correlation between Dice and relative lesion load was observed).

The pancreas’ anatomical flexibility and variability in shape, size, and location make it a more challenging organ for segmentation than the liver and the brain subcortical structures, which could explain the lower accuracy of the results by our atlas-based method for this organ. For comparison, recent works on pancreas segmentation applying CNNs to the first (TCIA) dataset report Dice scores as high as 0.83 [29, 31, 32].

Note that, in contrast to mainstream supervised segmentation methods that employ deformable registration or sophisticated trained neural networks, we compute the ELV map via a simple linear convolution operation on the (phase) image. Combining the signal-processing (ELV) and deep-learning (U-Net) approaches into a single neural network that receives both the image and the ELV map may improve the segmentation beyond each of the two approaches, which we propose as part of the future work.

## IV.Conclusions

We have introduced a new approach to supervised soft segmentation, which computes the expected label value (ELV) of a region of interest from an image using a training dataset of annotated atlases. The proposed method does not perform costly deformable registration, thereby also avoiding entrapment in local optima. We have evaluated the performance of our ELV computation technique in segmentation of the brain, the liver, and the pancreas. Future work consists of: using the ELV map to augment the input to a convolutional neural network beyond the image itself, expecting to increase the segmentation accuracy of the better-informed model; inherent inclusion of large rotation and scaling in ELV; and thorough evaluation with metrics other than Dice, such as region volume and surface distances.

## Supporting information

ELV soft-segmentation of the liver in the LiTS database.

Volumetric view of Figure 1.

## V. Appendices

### A. Inner Product as the Image Similarity Metric

The inner product of the new image *I* and the transformed atlas image *J* ° ***T***, which we have proposed as the image similarity metric in Eq. (3), is closely related to the sum-of-squared-differences (SSD) cost function that is commonly used in image registration:

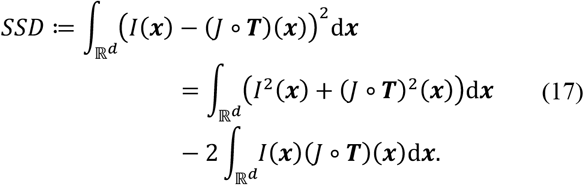

In order to establish an equivalence between maximizing our inner-product similarity function and minimizing SSD, it would seem necessary to include in Eq. (3) the term 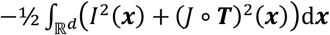, which is not necessarily constant with respect to ***T*** due to local volume changes in the transformation. The extra terms that such an addition would introduce in Pr (***T***|*I,J*)), of Eq. (2), however, can be seen to be independent of the global-translation component of ***T***. Then, since an integral with respect to ***T*** can be taken separately with respect to a global translation value and translation-free displacement fields, as in Eqs. (23)–(25), the extra terms in *E*(***y***) of Eq. (1) (resulting from the new translation-independent terms in Pr (***T***| *I,J*)) would be constant (independent of **y**), and therefore unnecessary in the computation of the ELV. Consequently, quantifying the similarity between two images as their inner product, as adopted here, corresponds to the common use of the SSD cost function in deformable image registration.

As mentioned in Section II.A, we use only the phase information of the images in Eq. (3), and measure the image similarity with the following inner product:

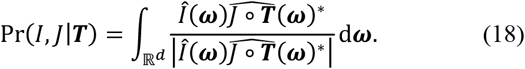

Using only the phase of the images, as in Eq. (18), is more suitable for the estimation of Pr (*I,J* |***T***), as it produces sharper probability distributions [16, 17]. To demonstrate this via an example, let us model the transformation as a simple translation, ***T*** (***x***)= ***x*** + Δ. The inner product therefore becomes the cross-correlation of the phase images, similar to Eq. (4), with Eq. (18) exhibiting the anticipated normality property, 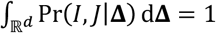 (although Pr (*I,J* |Δ) can occasionally become negative). Subsequently, in the simplistic case where *J* is a shifted version of, i.e. = *J* (***x***)=*I* (***x*** − Δ _**0**_), Eq. (18) will lead to Pr (*I,J* |Δ)= δ (Δ − Δ_**0**_), which is the exact desired distribution here.

Lastly, the inner product is zero for non-overlapping *I* and *J* ° ***T***, which is a crucial property for the image similarity metric in ELV computation.

### B. Incorporation of Deformation

In this appendix, we derive the ELV while accounting for deformations in the transformation (Section II.A.2). By combining Eqs. (1), (2), (3), and (9), the ELV at voxel ***y*** will be:

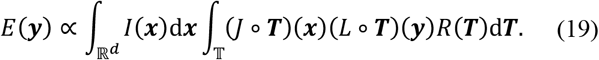

Since ***x*** and ***y*** are fixed in the inner integral, we make the change of variables ***T*** (z) = *S*(z – ***x***). Note that such a global shift will not change either the regularization, i.e.*R*(***T***)= *R*(*S*), or the domain of the inner integral, 𝕋 .Consequently:

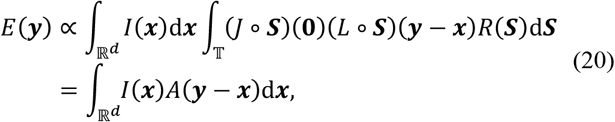

or:

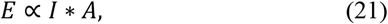

where we define the *key, A*, as:

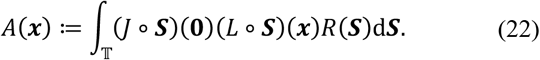

Next, we write the transformation as the sum of a global translation **Δ** ∈ ℝ^*d*^ and a deformation (displacement) field ***u*** ∈ *U*:

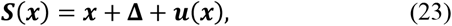

where 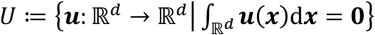 is the set of translation-free displacement fields. The regularity prior is now:

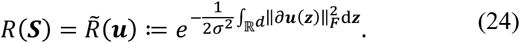

We combine the above three equations, and separate the integral over the space of all transformations into an integral over possible translation-free deformations and an integral over possible translations:

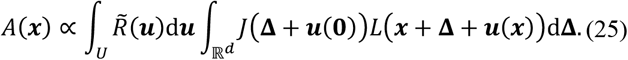

Note that this is a linear and invertible change of coordinates, hence *dS* ∝ Δ *d*u*d* (with the ratio independent of ***S***). With u and x being constant in the inner integral, we make the change of variables 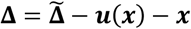:, leading to:

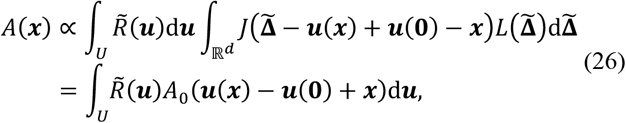

where *A*_**0**_ is the key for the translation-only case, introduced in Eq. (8):

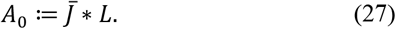

It can be verified that:

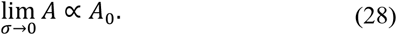

We now analytically estimate the key, *A*, as a function of *A*_**0**_ for *σ* > 0. Combining Eqs. (24) and (26) leads to:

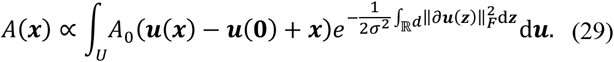

For simplicity, let us for now assume that ***x*** lies on the positive half of the first Cartesian coordinate axis, i.e., ***x*** = ***x****v*_1_, where *v*_1_ is the unit vector in the direction of the first axis, and ***x*** ≥ 0. We also define the line segment Q_***x***_ ≔ {***T***v_1_|0 ≤ ***T*** ≤ ***x***}. Accordingly:

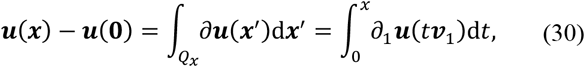

where ∂_1_***u*** is the partial derivative of ***u*** in the direction of ***v***_1_. Therefore:

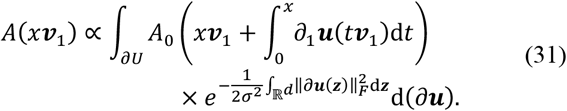

Note that we made further simplifying approximation by integrating over the space of the Jacobian of the deformation, ∂ U, instead of the space of the deformation, *U*, itself.^7^

In Eq. (31), the only values of ∂***u*** on which *A*_**0**_ depends are ∂_1_***u*** (***z***) for ***z*** ∈ *Q*_***x***_. Thus, we separate the integral into the product of three integrals, the first one being:

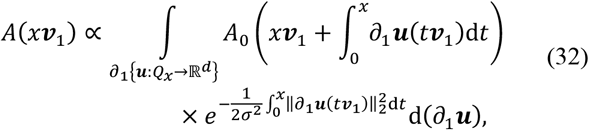

and the second and third integrals are:

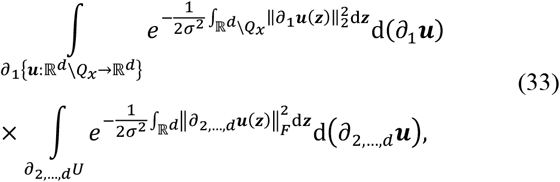

which are integrals of normal distributions and therefore constant, hence not included in the expression for *A*(***xv***_1_)in Eq. (32).

Calculation of *A* (***xv***_1_)can be made notationally easier by approximating the inner integrals in Eq. (32) as Riemann sums. We divide [0, ***x***] into equal intervals (*n* → ∞), with *d****T*** ^≈^ ***x***/ *N*, and define:

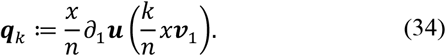

The integral is now approximated as:

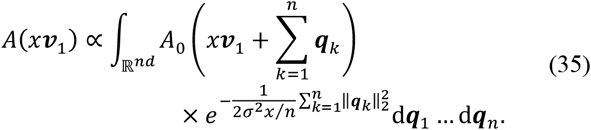

This is, in fact, *n* consecutive convolutions of *A*_**0**_ with a *d*-dimensional Gaussian,

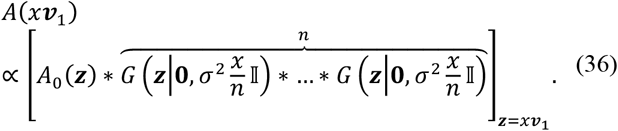

Given that convolution of *n* identical Gaussians results in a Gaussian with times the variance, we have:

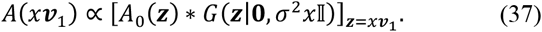

We now exploit the rotational invariance of the Gaussian in Eq. (37) and that of the Frobenius norm of the Jacobian in Eq. (29), to generalize Eq. (37) for any **x** ∈ ℝ^*d*^:

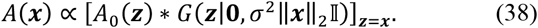

Equation (38) is indeed the update presented in Eq. (10). Despite our use of the convolution notation in Eq. (38), *A* is not computed via an actual convolution, because the co-variance matrix of the Gaussian kernel varies depending ***x*** on, where the result of the convolution is evaluated.

### C. Volume Threshold

To threshold the computed probability map of the organ, such as the ELV, we sort the values of the map and keep the top *v*^*∗*^ voxels, where the optimal *v*^*∗*^ needs to be determined. Assuming that the ground-truth label has *v*_*G*_ voxels, we define *L*(η)as the value of the ground-truth label at the top *v*^*Th*^ voxel, where *v* ≔ *ηv*_*G*_. An ideal probability map, whose top v_*G*_ voxels are the ground-truth label, is expected to produce the following boxcar function:

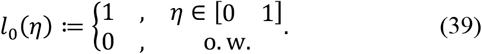

In practice, however, the transition to zero at *η* = 1 is less sharp due to inaccurately classified voxels, which we approximate with the following inverted sigmoid function:

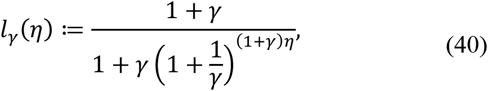

where *γ* is a nonnegative constant. Note that 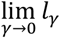 is the *L*_**0**_ defined in Eq. (39) for the ideal probability map, and that 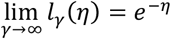. Furthermore, the normality of *L*_γ_, i.e. 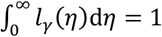, guarantees the expected property of 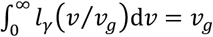.

Keeping the top *v* voxels results in a mask that overlaps with the ground-truth label with the following Dice similarity coefficient:

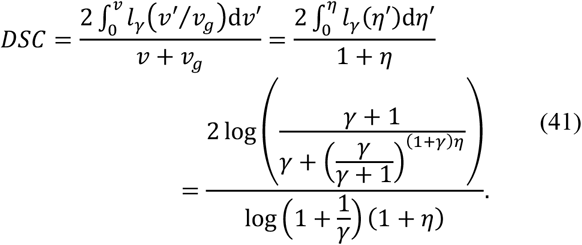

One can verify that, depending on the value of *γ*, the 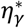 that maximizes the above Dice score ranges from 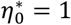 to 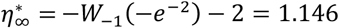, where W_k_ is the branch of the Lambert W function. Therefore, according to this model, the optimal number of top voxels of the probability map to keep (to maximize Dice) is 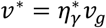. Choosing the nominal value of *γ* = 1 results in 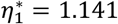, which led us to keep the top subset of voxels with a volume 14% larger than that of an average organ (Section II.C.1).^8^ Note that subsequent keeping of only the largest CCs in the resulting mask reduces the number of false-positive voxels, which, in our experiments, outweighed the newly introduced false-negative voxels, further increasing the Dice score.

## Footnotes

1 The gold-standard segmentation may also be a soft label, *L*: ℝ^*d*^ → [0,1].

2 We denote vector-valued variables in bold.

3 For data that is not too noisy, the organ size can also be estimated as the inflection point of the curve obtained by sorting the ELV map in descending order. Alternatively, the ELV map can be thresholded with a value optimized from the training data.

4 A Markov random field prior on the voxel labels could also be used to encourage spatial regularity [3, 19].

5 In our preliminary experiments, initial affine registration did not improve the results.

6 Resampling to a different resolution is only needed if the dimensions of the imaged object (in pixels) and their relative ratios substantially vary across subjects due to varying voxel size.

7 This change of variables (integrating with respect to *∂*u instead of u) is linear due to the linearity of the differential operator *A*, as well as invertible due to the translation-free constraint on u. We continue with the relaxing assumption that ∂u has independent elements. Nevertheless, for *d* ≥ 2, the variable set *A*u is redundant and has a larger dimension than u does, with elements that are interdependent given the linear relationship **∇** × *∂*u = **0**. As a result, for an exact solution, the integral must be taken with respect to an independent *subset* of the elements of *A*u that includes the (independent) set *∂*_1_u (Q_***x***_).

8 In our experiments, choosing *γ* = 1 consistently improved the results over γ = 0. The optimal value of *γ*, however, may vary depending on several factors, including the segmented organ, image modality and contrast, signal-to-noise ratio, etc.

